# Isolates from colonic spirochaetosis in humans show high genomic divergence and carry potential pathogenic features but are not detected by 16S amplicon sequencing using standard primers for the human microbiota

**DOI:** 10.1101/544502

**Authors:** Kaisa Thorell, Linn Inganäs, Annette Backhans, Lars Agréus, Åke Öst, Marjorie Walker, Nicholas J Talley, Lars Kjellström, Anna Andreasson, Lars Engstrand

## Abstract

Colonic spirochaetosis, diagnosed based on the striking appearance in histological sections, still has an obscure clinical relevance and only few bacterial isolates from this condition have been characterized to date. In a randomized, population-based study in Stockholm, Sweden, 745 healthy individuals underwent colonoscopy with biopsy sampling. In these individuals, 17 (2.3 %) had colonic spirochaetosis, which was associated with eosinophilic infiltration and a three-fold increased risk for irritable bowel syndrome (IBS). We aimed to culture the bacteria and perform whole genome sequencing of the isolates from this unique representative population sample. From 14 out of 17 individuals with spirochaetosis we successfully isolated, cultured and performed whole genome sequencing of in total 17 isolates including the *Brachyspira aalborgi* type strain 513A^T^. Also, 16S analysis of the mucosa-associated microbiota was performed in the cases and non-spirochaetosis controls.

This is the first report of whole genome analysis of clinical isolates from individuals with colonic spirochaetosis. We found one isolate to be of the species *Brachyspira pilosicoli* and all remaining isolates were of the species *Brachyspira aalborgi*. Besides displaying extensive genetic heterogeneity, the isolates harboured several mucin-degrading enzymes and other virulence-associated genes that could confer a pathogenic potential in the human colon. We also showed that 16S amplicon sequencing using standard primers for human microbiota studies fail to detect *Brachyspira* due to primer incompatibility. This failure to detect colonic spirochaetosis should be taken into consideration in project design and interpretation of gastrointestinal tract microbiota in population-based and clinical settings.

## Introduction

Colonic spirochaetosis (CS) was first described in 1967, when spirochaetes adherent to the surface of colonic epithelium were identified by electron microscopy of rectal biopsies in a seminal paper by Harland and Lee who coined the term intestinal spirochaetosis^1^. The characteristic histological appearance of the bacteria has been considered to be pathognomonic for diagnosis, but the biology and origin of intestinal spirochaetes in humans is still poorly understood.

The nosology for spirochaetes in the human intestine has varied from *Borrelia* and *Serpulina*, to the present classification *Brachyspira*, and spirochaetal bacteria have been identified in the intestines of several animals including monkeys, dogs, chickens, rodents and pigs ^2^. In animals with colonic spirochaetosis, a spectrum of medical conditions is well described. While rodents present with asymptomatic excretion, in swine the spirochaetes can cause pathologic changes leading to diarrhoea, malnutrition, and declining growth rates, resulting in high economic losses ^3^. Primates occupy a rather intermediate position, where even though spirochaetal organisms can colonize the colonic mucosa, the animals rarely present with enteric symptoms ^4^. In humans, at least two spirochaete species, *Brachyspira pilosicoli* and *Brachyspira aalborgi*, are associated with spirochaetosis ^5^, and rare findings of concomitant infection by *B. aalborgi* and *B. pilosicoli* have been described ^6,7^ These spirochaetes are both fastidious and rather slow-growing anaerobes ^8^, and successful culture of spirochaetes from stool or biopsy material is rarely reported in human studies. However, the type species of the genus *Brachyspira* (*B. aalborgi*) was originally isolated from a patient with diarrhoea ^9^ and more recently another strain isolated from a colonic biopsy in a patient with blood and mucus in stool was reported ^10^. However, no *B. aalborgi* and only four *B. pilosicoli* isolates have been genomically characterized on the whole genome level to date ^11^.

Even though there has been progress in detecting and identifying spirochaetosis in humans, studies based on histopathological diagnosis without detailed symptom correlation has left the question as to whether it represents a disease process unanswered ^12,13^. However, more recent studies have shown an association between spirochaetosis and the irritable bowel syndrome (IBS), and identified a unique colonic pathology characterized by increased eosinophils in this condition ^14,15^. As IBS affects one in 10 people worldwide, impairs quality of life and is highly costly, the observation that colonic spirochaetes are associated with IBS-diarrhoea is of major interest ^16^.

In a unique representative random population sample that underwent a study colonoscopy in Sweden, we aimed to culture and perform whole genome sequencing of colonic spirochaetosis isolates obtained from colonic biopsies. To characterize isolates from human spirochaetosis, we in this study cultured spirochaetal isolates from individuals with microscopically determined spirochaetosis from within the randomized, population-based colonoscopy study PopCol that was performed in Stockholm, Sweden in 2001-2006 ^17^. A total of 745 healthy adults underwent colonoscopy with biopsy sampling of four sites of the colon and of the terminal ileum. Out of these individuals, 17 presented with spirochaetosis, an observation that was shown to be associated with eosinophilic infiltration in the tissue and a three-fold increased risk for IBS ^14^. Since the spirochaetes were found solely in the colon (and not in the terminal ileum), we will hereafter use the term colonic spirochaetosis (CS) ^3^. To further characterize the spirochaete bacteria and their effect on the colonic microbiota, we performed whole-genome sequencing of the bacterial isolates together with type strain *513A^T^* [ATCC 43994] ^9^, and reference strain W1 ^10^, as well as performed 16S amplicon sequencing for microbiota profiling of colonic biopsies in the individuals with colonic spirochaetosis and unaffected controls. We here present the first whole-genome sequences of isolates from human colonic spirochaetosis, which display extensive genetic heterogeneity also between members of the same species.

## Materials and methods

### Population

The population-based colonoscopy (PopCol) study has been described in detail elsewhere ^17^. Briefly, a random sample of the adult population in two adjacent parishes in Södermalm, Stockholm, were sent a validated abdominal symptom questionnaire (the ASQ) ^18^. Of 3556 recipients, 2293 responded to the questionnaire and of these, 1643 persons were reached by phone with an invitation to an interview with a colonoscopist and a subsequent colonoscopy. A total of 1244 participated in the interview of which 745 also underwent colonoscopy. The subjects who underwent colonoscopy were similar to the total population enrolled ^17^. In 17 of the individuals spirochaetes were observed in Haematoxylin and Eosin-stained tissue sections, which was subsequently confirmed by immunohistochemistry using a polyclonal rabbit antiserum ^10,14^. The study was approved by the local Ethics committee (No 394/01, Forskningskommité Syd), and all participants gave their informed consent. Irritable bowel syndrome (IBS) and bowel habit subtypes (diarrhoea, mixed diarrhoea and constipation, and unclassifiable) was defined by applying Rome III criteria ^19^, and has been described in detail in Walker et al. 2015 ^14^.

### Collection of biopsies

The colonoscopies were performed by 7 experienced endoscopists. Biopsies were taken at 4 levels in the colon (caecum, transverse, sigmoid and rectum) and in terminal ileum. For more details of biopsy collection, see Kjellström et al. ^17^.

For both the isolation of bacteria and the 16S amplicon sequencing, sigmoid colon biopsies were used. For the microbiota analysis four age- and sex matched controls without spirochaetosis per CS case were randomly selected from the participants with normal pathology (both macroscopic and microscopic) from the PopCol study.

### Isolation and biochemical characterisation of spirochaetes

Biopsies from the 17 individuals with confirmed spirochetosis were homogenised in freezing media and 50 μl was cultivated anaerobically on selective medium for spirochaete isolation and subculturing as previously described ^10^. Pure growth of spirochaetal bacteria was confirmed with a phase-contrast microscope. From three of the individuals we could not isolate any spirochaetes by our methods. From one individual two isolates were selected since they were displaying different morphology on the plate. *B. aalborgi* type strain *513A^T^* [ATCC 43994 / NCTC 11492] ^9^, and the Swedish clinical *B. aalborgi* isolate W1 ^10^, from a strain collection at the National Veterinary Institute, Uppsala, were also cultured and used for comparison. See Table 1 for a summary of the individuals and isolates.

**Table 1.**
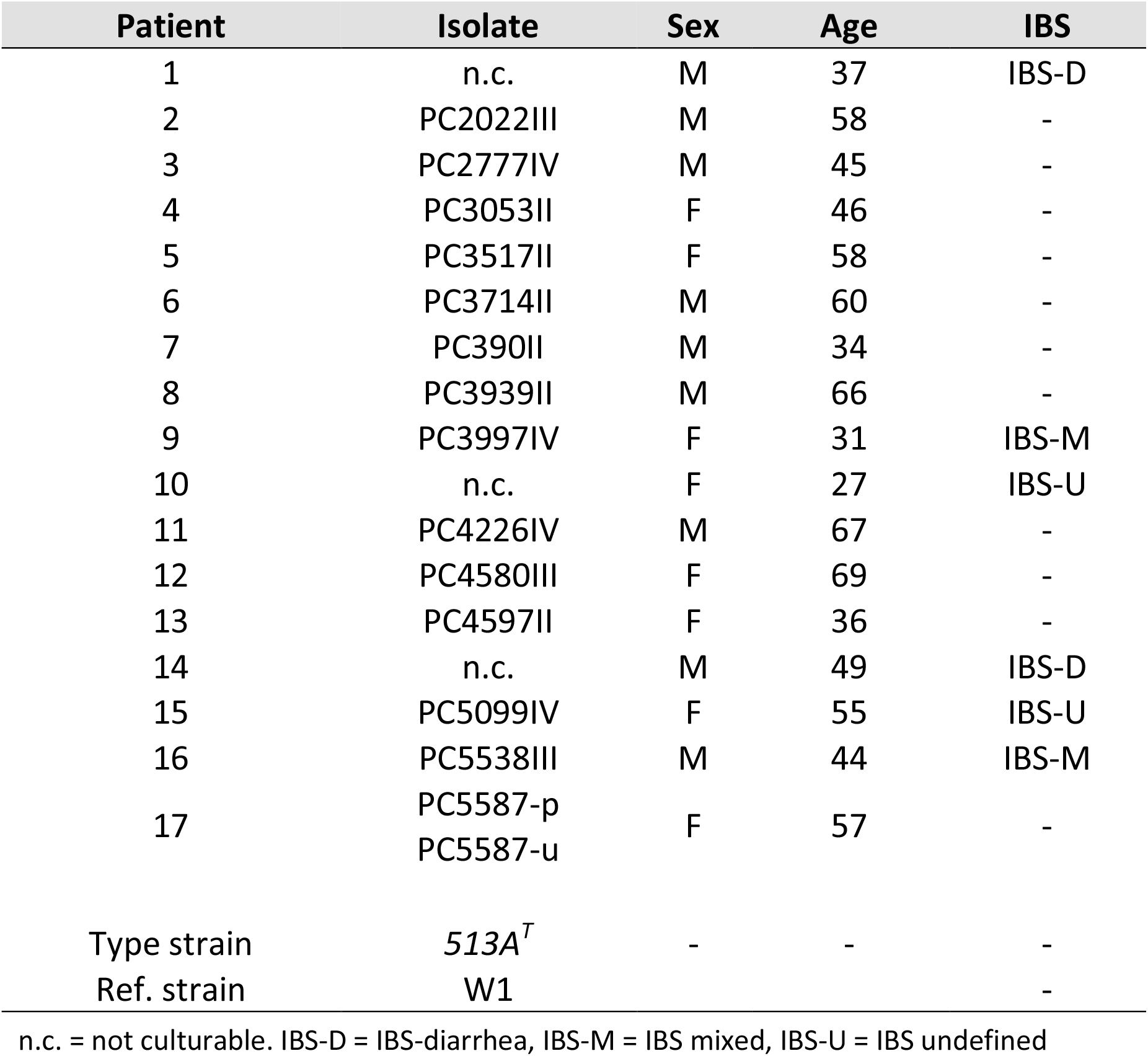
Subject and isolate information.

| Patient | Isolate | Sex | Age | IBS |
| --- | --- | --- | --- | --- |
| 1 | n.c. | M | 37 | IBS-D |
| 2 | PC2022III | M | 58 | - |
| 3 | PC2777IV | M | 45 | - |
| 4 | PC3053II | F | 46 | - |
| 5 | PC3517II | F | 58 | - |
| 6 | PC3714II | M | 60 | - |
| 7 | PC390II | M | 34 | - |
| 8 | PC3939II | M | 66 | - |
| 9 | PC3997IV | F | 31 | IBS-M |
| 10 | n.c. | F | 27 | IBS-U |
| 11 | PC4226IV | M | 67 | - |
| 12 | PC4580III | F | 69 | - |
| 13 | PC4597II | F | 36 | - |
| 14 | n.c. | M | 49 | IBS-D |
| 15 | PC5099IV | F | 55 | IBS-U |
| 16 | PC5538III | M | 44 | IBS-M |
| 17 | PC5587-p<br>PC5587-u | F | 57 | - |
| Type strain | 513A<sup>T</sup> | - | - | - |
| Ref. strain | W1 |  |  | - |

n.c. = not culturable. IBS-D = IBS-diarrhea, IBS-M = IBS mixed, IBS-U = IBS undefined

The spirochaetes were tested for indole production, their ability to hydrolyse sodium hippurate, beta-haemolysis capacity, and cellular α-galactosidase and β-glucosidase activity, as has been described previously ^10^

### DNA extraction and sequencing library preparation of spirochaetes

The spirochaetes were harvested and DNA was extracted from the pellet using the DNeasy Blood & Tissue kit (Qiagen, Hilden, Germany) with proteinase K treatment at 56 C° for enhanced lysis.

Library preparation was performed using the TruSeq Nano DNA library preparation kit (Illumina) using the 550 bp insert size protocol and the libraries were subsequently sequenced on the MiSeq platform, v2 chemistry, 300 bp paired end reads. The average coverage of the genomes was >300-fold and the average insert size 900 bp. For more detailed sequencing and assembly statistics, see Table S1.

## Bioinformatics analysis of whole genome sequences

### De novo genome assembly and annotation

Reads were quality assessed and trimmed using bbduk (bbmap v. 37.77) ^20^ and *de novo* assembled using SPAdes v 3.11.1 ^21^ assembly, using the --careful option and kmer options 21, 33, 55, 77, 99, and 127, and the assemblies were filtered to remove contigs of low coverage or a length under 500 bp. The draft genome of strain *513A^T^* was ordered using the Mauve ^22^ Order Contig function to the incomplete draft of *Brachyspira aalborgi 513A* available at the MetaHIT consortium repository (https://www.sanger.ac.uk/resources/downloads/bacteria/metahit/) and the other draft genomes were subsequently ordered to the *513A^T^* draft. The ordered drafts were annotated using the *prokka* annotation pipeline v 1.12 ^23^.

### Species Circumscription and Phylogeny

To investigate the species designation of the CS isolates, phylogenetic analysis of 16S rRNA sequences was performed by comparing the 16S sequences of the human CS strains to the Ref and RefNR *Brachyspira* 16S sequences retrieved from the SILVA database (https://www.arb-silva.de/). The sequences were aligned using muscle ^28^ and a Maximum Likelihood tree was constructed using PhyML ^29^, both software integrated in Seaview v. 4.5.4^30^.

The NADH oxidase (*nox*) gene has often been used for phylogenetic comparison of spirochaetes ^31^ and was analysed by combining publically available *Brachyspira nox* sequences from GenBank (Table S3) and the *nox* sequences from our newly sequenced strains. The nucleotide sequences were aligned using muscle and a phylogenetic tree was reconstructed using PhyML as described above.

To study the genetic variability between the CS strains and hitherto described genomes within the *Brachyspira* genus listed in Table S1, an average nucleotide identity (ANI) analysis was performed using MUMmer alignment ^32^ implemented in the *pyani* software v. 0.2.7 (https://github.com/widdowquinn/pyani).

*B. sp*. CAG-484 (<u>GCA_000431315.1</u>), a metagenomic assembly from the metaHIT consortium was excluded from these analyses since it was highly divergent from the others and seemed to be misclassified to the *Brachyspira* genus.

Digital DNA-DNA hybridization (dDDH) analysis was performed using the Genome-to-Genome Distance Calculator (GGDC) web tool v. 2.1 ^33^.

To compare the genetic content of the isolates, the Roary pan-genome pipeline ^34^ was used, with an identity cut-off of 70% on protein level. This analysis was performed both including all the above-mentioned *Brachyspira* genome references and then separately for the *B. pilosicoli* genomes, and only including the *B. aalborgi* isolates.

A more detailed analysis of strains associated with human colonic spirochaetosis was also performed by retrieving the 16S sequences described in Mikosza et al ^35^, Pettersson et al. ^36^ and Westerman et al ^37^ (Table S2). The sequences were aligned and a PhyML tree was constructed as described above.

### Comparative genomics and functional genomics analysis

To further investigate the functional characteristics of the CS genomes, we further scrutinized the Roary pan-genome data by classifying the representative sequences of each core and accessory gene cluster according to KEGG orthology using the BlastKOALA online tool ^38^. We also classified all *B. aalborgi* reference strain *513A^T^* genes into Clusters of Orthologous Groups (COG) categories ^39^ to facilitate functional interpretation. To further functionally characterise the genomes we applied the macsyfinder *TXSScan* tool ^40^ to identify bacterial secretion systems (T1SS-T6SS, T9SS, flagella, type IV pili and Tad pili) and the prophage prediction tool ProPhet ^41^.

## Colonic microbiota composition in individuals with CS compared to controls

### DNA extraction

The homogenised sigmoid colon biopsies were transferred from their original freezing media into DNA/RNA Shield lysis buffer prior to extraction with ZymoBIOMICS (Zymo Research Corp, Irvine, CA).

The biopsies were submitted to bead beating with Matrix E (MP Biomedicals, Santa Ana, CA, USA) in a 96 FastPrep shaker (MP Biomedicals, Santa Ana, CA, USA) for 2-6 minutes (the beating proceeds until the sample is visually homogeneous) and samples were then spun down to remove beads from the solution. The supernatants were then incubated in lysozyme buffer (20 mM Tris-CL, 2 mM sodium-EDTA, lysozyme to 100 g/mL; Sigma, St. Louis, MO, USA) at 37°C for 45’ to 1h at 1000 RPM. Following this, samples were again spun down and the supernatant transferred to a new plate, to eliminate larger particles, and then incubated with proteinase K at 55°C at 250 RPM for 30 minutes. Finally, the samples were cleaned through several washing and magnetic bead pelleting steps according to the instructions of the manufacturer (Genomic DNA MagPrep kit, Zymo Research Corp, Irvine, CA, USA) and the DNA eluted from the magnetic beads with 70 μL of Elution Buffer (10 mM Tris-Cl, pH 8.5; Qiagen, Venlo, Netherlands). The DNA extraction was automated on the FreedomEVO robot (TECAN Trading AG, Männendorf, Switzerland).

A blank negative control and a positive mock control (ZymoBIOMICS Mock Community Standard, Zymo Research Corp, Irvine, CA, USA) were included in each extraction round.

### DNA amplification and sequencing

Prior to amplification samples were normalized and a total 170 ng DNA was used to amplify the V3-V4 region of the 16S rRNA gene using primer pair 341F/805R ^42^.

For the 1-step PCR procedure, amplification was carried out by a high fidelity proofreading polymerase for a total of 25 cycles. For amplification of the sequencing libraries, forward primer 5’-CAAGCAGAAGACGGCATACGAGAT-N_8_-GTCTCGTGGGCTCGGAGATGTGTATAAGAGACAGGACTACHVGGGTATCTAATCC-3’ and reverse primer 5’AATGATACGGCGACCACCGAGATC-N_8_-TCGTCGGCAGCGTCAGATGTGTATAAGAGACAGCCTACGGGNGGCWGCAG-3’, where N_8_ represents an identifying 8-mer (barcode) and the last 21 and 19 bases in each construct are the sequence specific forward and reverse primers, respectively, were used.

Samples were then pooled to equimolar amounts and sequenced in parallel to whole bacterial genomes on the MiSeq instrument (Illumina Inc., San Diego, CA, USA). All controls from the extraction phase, as well as a negative (blank) PCR control were also prepared and sequenced.

### Sequence correction and taxonomic assignment

The processing of 16S amplicon reads was performed as described previously ^43^. Briefly, Cutadapt ^44^ was used to eliminate all sequences not containing the amplification primers, remove the primer sequences, all bases with a Phred score below 15 and all reads with less than 120 bp left after trimming. The resulting reads were merged with usearch v.9.0.2132 ^45^ and reads failing to merge or producing merging products shorter than 380 bp, longer than 520 bp or with more than three expected errors were discarded. All unique full-length samples occurring at a frequency higher than 10^−6^ in the dataset were submitted to unoise ^46^. All the merged reads were then mapped back to the accepted centroids and assigned to the OTU with the highest identity, at a minimum of 98%. In case of equally high identity matches, the most abundant centroid is selected. Taxonomy was assigned using SINA v.1.2.13 ^47^. Scripts for performing this analysis are available at https://github.com/ctmrbio/Amplicon_workflows.

### Statistical analysis

For statistical analyses, all samples with fewer than 5000 total Operational Taxonomic Units (OTU) counts were discarded, as were OTUs not present in at least 3 samples. The resulting filtered OTU counts were parsed to group counts on the different taxonomical levels and visualised using the R software packages ggplot ^48^ and pheatmap ^49^. Shannon and Chao1 alpha diversity statistics and Bray-Curtis similarity were calculated using the vegan R package ^50^. Differences in diversity statistics between Spirochetosis cases and controls were tested using student’s t-test.

### Verification of 16S primer performance

Due to the low total counts of reads from the biopsy amplicon sequencing assigned to the *Brachyspira* genus we performed a control experiment to verify the amplification performance of the 341F/805R primer combination on mixes of bacterial DNA. The mixes were prepared of 75% ZymoBIOMICS™ Microbial Community DNA Standard and 25% DNA from five of the different *Brachyspira* isolates, selected from different clades of the 16S rRNA phylogenetic tree (Figure 2), namely *B. aalborgi 513A^T^*, W1, PC3939II and PC3714II, and *B. pilosicoli* PC5538III. The standard is a defined mixture of 5 Gram-positive and 3 Gram-negative bacteria plus 2 yeast species with wide GC range (15%-85%) for evaluation and optimisation of microbiomics workflows. A pure ZymoBIOMICS™ Microbial Community DNA standard sample and a blank sample were also included as positive and negative PCR control.

### Analysis of Human Microbiome Project data

To search for the presence of spirochaetes in the publically available microbiota data from the Human Microbiome Project (HMP) we downloaded 16S amplicon data from 324 stool samples and metagenomic assemblies from 179 stool samples (Table S5). The 16S samples were analysed with the methods described above while the assembled shotgun metagenome data was screened for the presence of spirochetes using MASH ^51^. Briefly, the colonic spirochetosis *Brachyspira aalborgi* and *pilosicoli* genomes previously described in this study were combined and a MASH sketch was created from these, to which all the downloaded assemblies were compared. The assemblies showing signs to contain spirochaete sequences were aligned to the combined genomes using blastn ^52^ and the contigs having hits with an e-value of lower than e^−9^ were extracted for further investigation.

## Results

### Isolation of Spirochaetes

In the present study, spirochaetes were successfully isolated from frozen biopsies from 14 out of the 17 individuals (Table 1). All three individuals from whom we were not able to retrieve viable bacteria were in the IBS group. From one of the individuals, colonies of different morphologies were identified and two isolates were propagated, PC5587-p and PC5587-u. In addition to the clinical isolates, *B. aalborgi* type strain *513A^T^* [ATCC 43994/NCTC 11492] ^9^, and the Swedish *B. aalborgi* reference isolate W1 ^10^ were sequenced.

### Morphological and biochemical characterization of the isolates

Originally identified in Haematoxylin and Eosin-stained sections and confirmed with immunohistochemistry we could observe the spirochaetes to completely cover the colonic epithelial membranes in all individuals. This could also be visualized using Warthin-Starry silver stain and representative pictures of the histological manifestation are shown in Figure 1. The spirochetes were present at all the colon levels studied except for in rectal biopsies where we could not detect the spirochaetes in 4 out of 17 individuals. In the tissue sections the bacteria were of similar morphology in all individuals with medium-long spiral shaped bacteria seemingly adherent to the colon mucosa. In culture, the growth on the fastidious anaerobic agar plates was mostly weak with small colonies and occasional swarming. The isolates, in contrast to what was observed in the tissue sections, showed a considerable variation in phenotype, both in size, growth rate, cell size, and biochemical reactions (Table 2).

**Figure 1.**
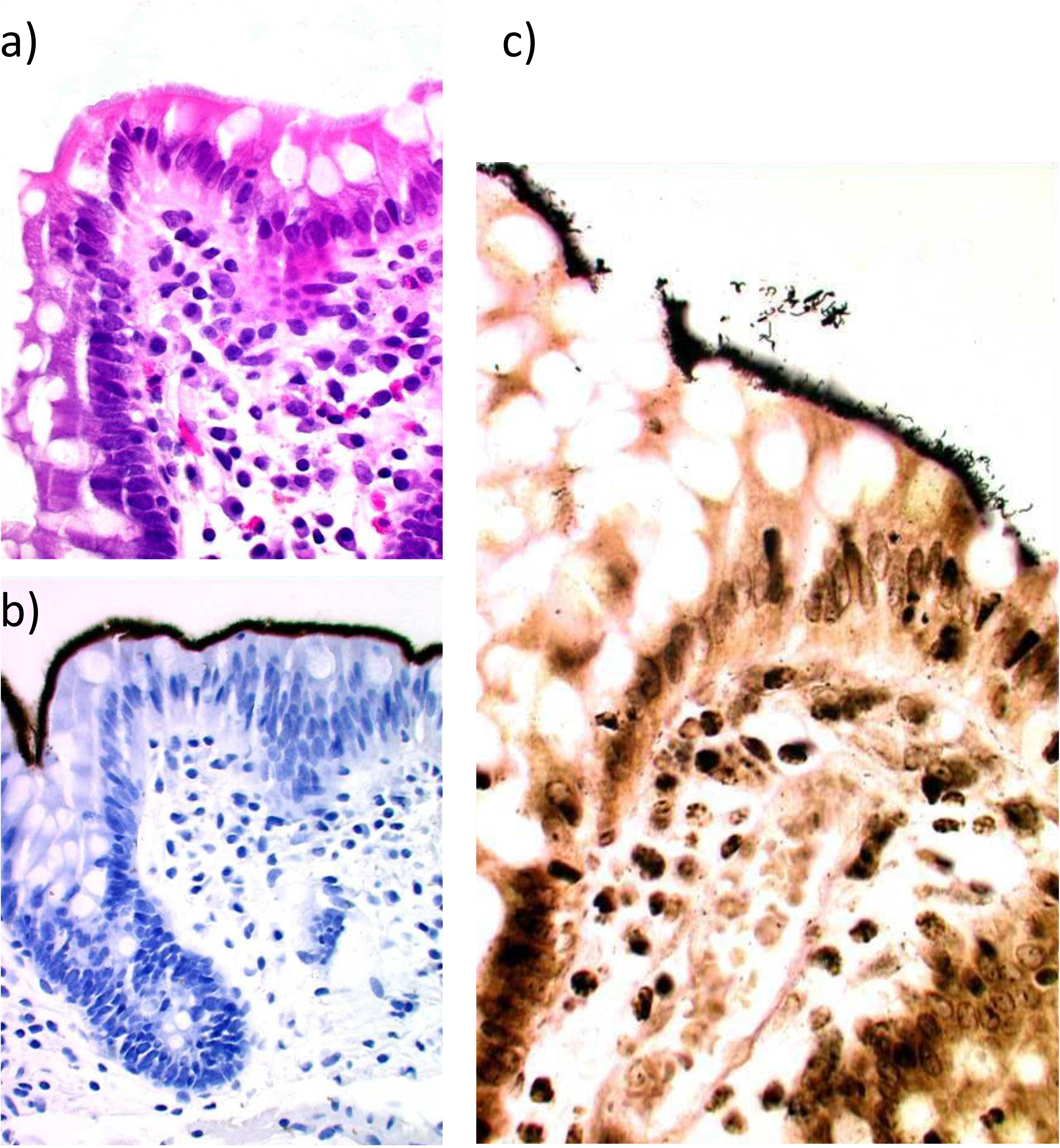
Histological sections showing, in the same formalin-fixed, paraffin embedded biopsy, representative pictures of the spirochaetosis using A) Haematoxylin and Eosin staining, B) Immunohistochemistry using polyclonal spirochaete-specific antibodies, C) Warthin-Starry Silver staining.

**Table 2.** Biochemical analyses.

|  |  | Biochemical test |  |  |  |  |
| --- | --- | --- | --- | --- | --- | --- |
| | Isolate | Haemolysis | Indole production | Hippurate hydrolysis | α-gal<sup>a</sup> | β-glu<sup>b</sup> |
| Clinical Isolates | PC4226IV | Weak | - | - | - | + |
|  | PC3053II | None | + | - | - | - |
|  | PC4580III | None | - | - | - | - |
|  | PC3997IV | Weak | - | - | - | - |
|  | PC2777IV | Weak | - | - | - | - |
|  | PC3939II | Weak | + | - | - | + |
|  | PC3517II | Weak | + | - | - | + |
|  | PC3714II | None | - | - | - | + |
|  | PC5099IV | None | - | - | - | - |
|  | PC5538III | None | + | - | - | - |
|  | PC4597II | None | - | - | - | (+) |
|  | PC390II | None | - | - | - | (+) |
|  | PC2022II | None | - | (+) | - | - |
|  | PC5587II-p | None | - | - | - | - |
| Type strain | 513A<sup>T</sup> | None | - | - | - | - |
| Ref. strain | W1 | None | - | (+) | - | - |

<sup>a</sup> $\alpha$ -gal = alpha-galactosidase activity.
<sup>b</sup> $\beta$ -glu = beta-glucosidase activity.

Out of the 14 spirochaetal isolates that were investigated, one isolate shared reaction pattern with the reference strain W1 and five isolates shared reaction patterns with the type strain *513A^T^* (Table 2). Eight strains showed different reaction patterns including four indole positive isolates. Two isolates, including the reference strain W1 had a positive but weak hippurate hydrolysis. When compared to the results from the phylogenetic analysis there was no obvious correlation.

### Genome assembly and phylogenetic classification

Whole genome *de novo* assembly of the 17 isolates revealed that the bacteria had genomes of approximately the same size, on average 2.67 Mbp with the smallest being *513A^T^* with a genome of 2.50 Mbp (Table 3). The PC5538III culture turned out to contain two isolates that could be separated in the assembly based on coverage over the contigs. These were termed PC5538III-hc and PC5538III-lc for high- and low coverage, respectively. The GC content varied between 27.6% in PC5538III-hc to 28.3%. Most of the genomes assembled nicely with an average of 31 (8-113) contigs per genome. For more detailed assembly statistics, see Table S4.

**Table 3.**
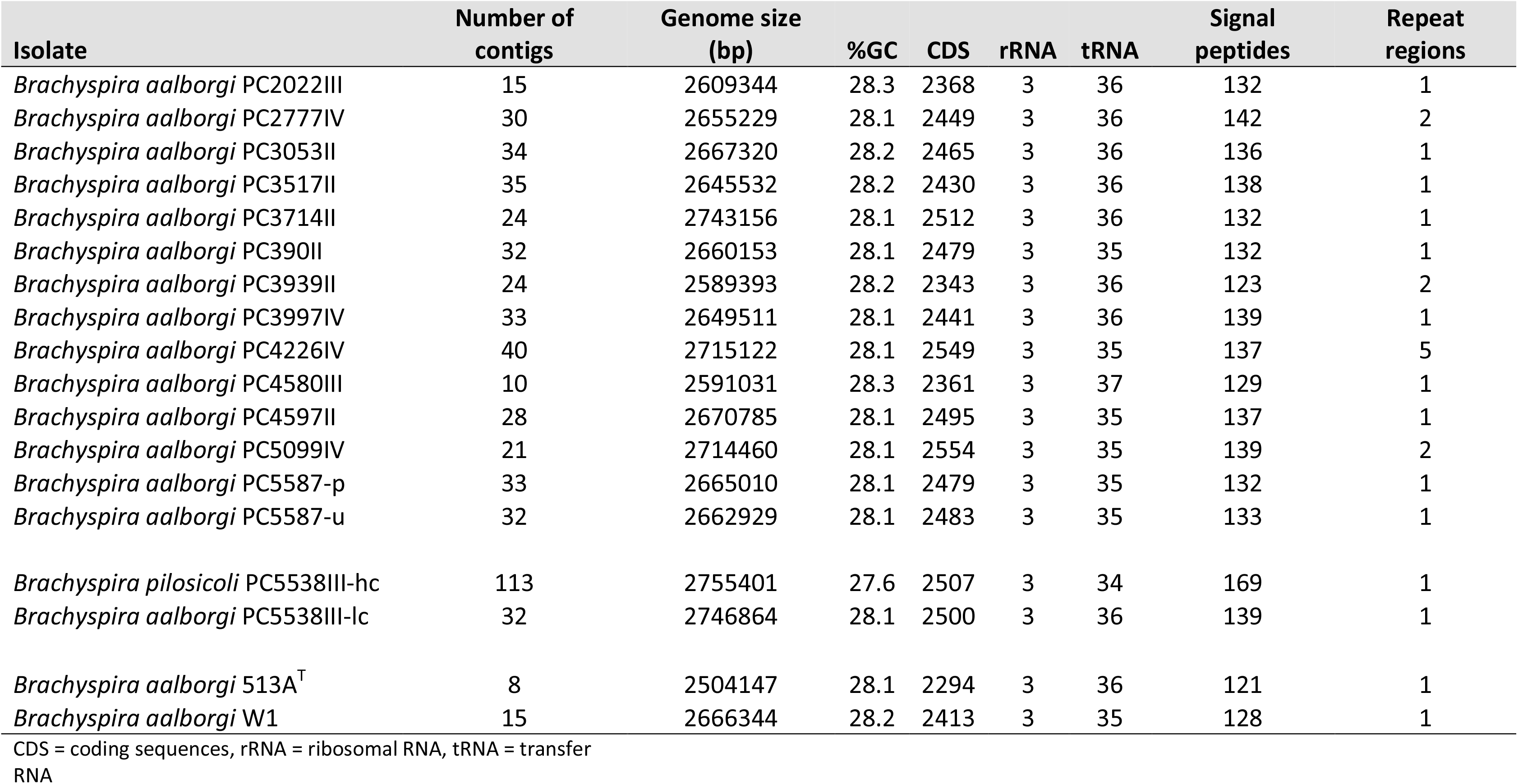
Genome characteristics.

| Isolate | Number of contigs | Genome size (bp) | %GC | CDS | rRNA | tRNA | Signal peptides | Repeat regions |
| --- | --- | --- | --- | --- | --- | --- | --- | --- |
| *Brachyspira aalborgi* PC2022III | 15 | 2609344 | 28.3 | 2368 | 3 | 36 | 132 | 1 |
| *Brachyspira aalborgi* PC2777IV | 30 | 2655229 | 28.1 | 2449 | 3 | 36 | 142 | 2 |
| *Brachyspira aalborgi* PC3053II | 34 | 2667320 | 28.2 | 2465 | 3 | 36 | 136 | 1 |
| *Brachyspira aalborgi* PC3517II | 35 | 2645532 | 28.2 | 2430 | 3 | 36 | 138 | 1 |
| *Brachyspira aalborgi* PC3714II | 24 | 2743156 | 28.1 | 2512 | 3 | 36 | 132 | 1 |
| *Brachyspira aalborgi* PC390II | 32 | 2660153 | 28.1 | 2479 | 3 | 35 | 132 | 1 |
| *Brachyspira aalborgi* PC3939II | 24 | 2589393 | 28.2 | 2343 | 3 | 36 | 123 | 2 |
| *Brachyspira aalborgi* PC3997IV | 33 | 2649511 | 28.1 | 2441 | 3 | 36 | 139 | 1 |
| *Brachyspira aalborgi* PC4226IV | 40 | 2715122 | 28.1 | 2549 | 3 | 35 | 137 | 5 |
| *Brachyspira aalborgi* PC4580III | 10 | 2591031 | 28.3 | 2361 | 3 | 37 | 129 | 1 |
| *Brachyspira aalborgi* PC4597II | 28 | 2670785 | 28.1 | 2495 | 3 | 35 | 137 | 1 |
| *Brachyspira aalborgi* PC5099IV | 21 | 2714460 | 28.1 | 2554 | 3 | 35 | 139 | 2 |
| *Brachyspira aalborgi* PC5587-p | 33 | 2665010 | 28.1 | 2479 | 3 | 35 | 132 | 1 |
| *Brachyspira aalborgi* PC5587-u | 32 | 2662929 | 28.1 | 2483 | 3 | 35 | 133 | 1 |
| *Brachyspira pilosicoli* PC5538III-hc | 113 | 2755401 | 27.6 | 2507 | 3 | 34 | 169 | 1 |
| *Brachyspira aalborgi* PC5538III-lc | 32 | 2746864 | 28.1 | 2500 | 3 | 36 | 139 | 1 |
| *Brachyspira aalborgi* 513A<sup>T</sup> | 8 | 2504147 | 28.1 | 2294 | 3 | 36 | 121 | 1 |
| *Brachyspira aalborgi* W1 | 15 | 2666344 | 28.2 | 2413 | 3 | 35 | 128 | 1 |

CDS = coding sequences, rRNA = ribosomal RNA, tRNA = transfer RNA

To identify the species of the isolates, ribosomal RNA sequences were extracted and searched against the SILVA rRNA database, identifying PC5538III-hc to be *Brachyspira pilosicoli* and all other isolates to belong to *Brachyspira aalborgi* (Figure 2A). Another approach commonly used for the phylogenetic comparison between *Brachyspira* strains is to use the NADPH oxidase (*nox*) gene ^31^. We therefore combined the publicly available *Brachyspira nox* sequences (Table S3) with the *nox* sequences from our strains. The phylogenetic tree can be seen in Figure 2B and confirms the classification of all but one of the strains to *B. aalborgi*, grouping together with the previously sequenced *nox* gene of *B. aalborgi ATCC 43994 [513A^T^]* and the last, PC5538III-hc to *B. pilosicoli*.

**Figure 2.**
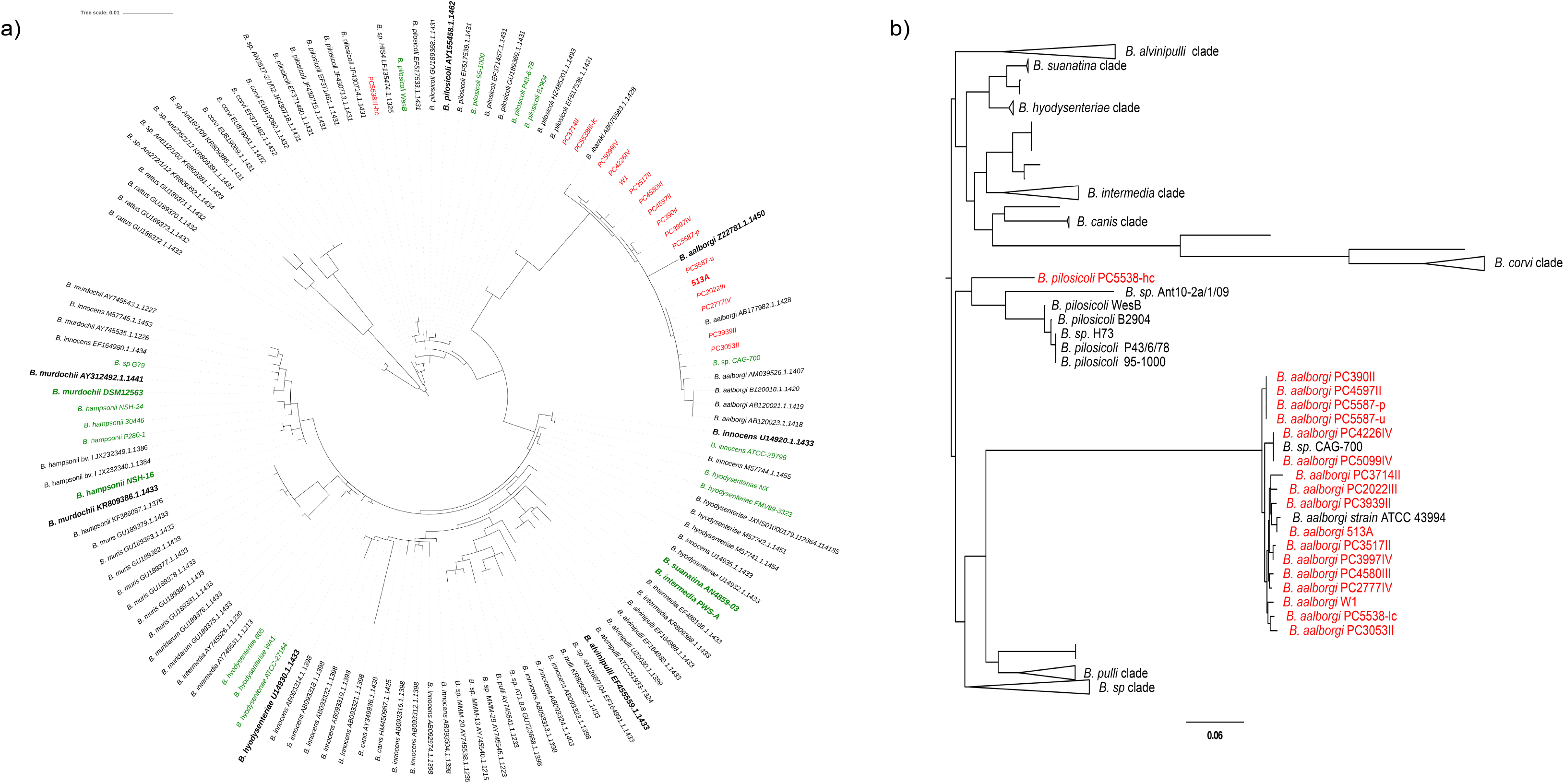
Phylogenetic trees based on A) the 16S ribosomal RNA sequences, and B) the NADH oxidase (NOX) protein sequence respectively. The isolates marked in red were sequenced within this study. The sequences marked in green come from the whole-genome sequences described in Table S1. The taxa in bold are the type strains for their respective species. In panel B clades containing only sequences from one species are collapsed for better readability.

To investigate the diversity between the *Brachyspira* isolates we also compared the genomes to publicly available whole genome sequences within the *Brachyspira* genus, both using average nucleotide identity (Fig. S1A, Table S6, digital DNA-DNA hybridization (dDDH) (Fig. S1B), and based on gene content in a pan-genome analysis (Fig. S1C), confirming the species identification. Also, we could confirm the previously unclassified *Brachyspira sp. CAG-700*, derived from a human gut metagenome ^53^ to belong to the *B. aalborgi* species and it was therefore included in further comparative analyses of the species.

Since there are no previous studies reporting whole-genome sequences of isolates from colonic spirochaetosis we made a comparison on 16S rRNA level with published rRNA sequences from three previous studies on human intestinal spirochaetosis; Pettersson et al. 2000 ^36^, Mikosza et al. 2004 ^35^ and Westerman et al. 2012 ^37^, (Table S2). The CS isolates were shown to belong to different clusters of the *B. aalborgi* tree (Fig. 3). None of the isolates fell within the clade proposed to be called *Brachyspira hominis* ^37^, while isolate PC3714II grouped away from the others, forming its own branch between *B. aalborgi* and *B. pilosicoli*. Also, the analysis showed that all our isolates clustered with previously described cluster 1 isolates ^35,36^.

**Figure 3.**
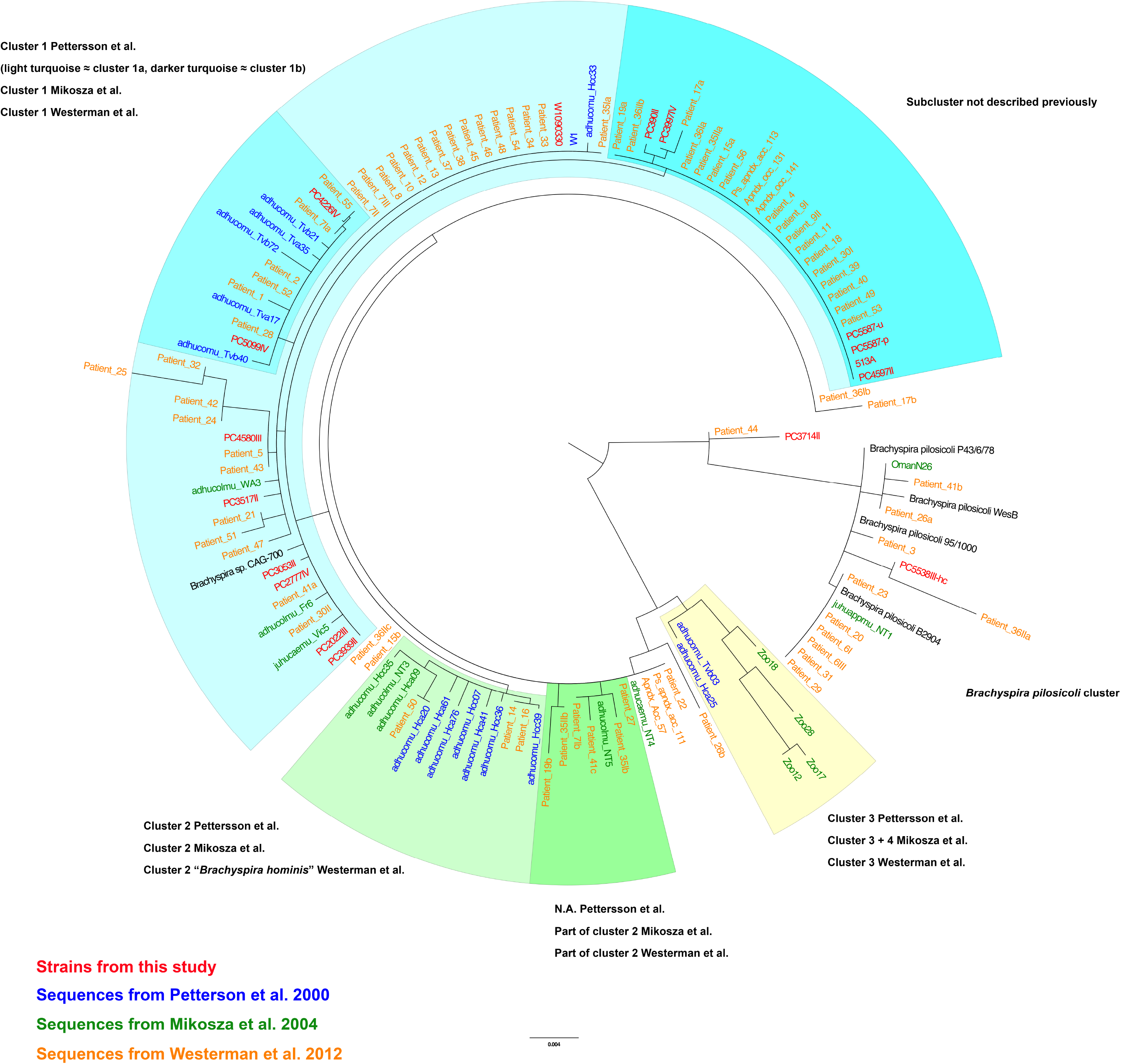
Maximum likelihood tree of 16S rRNA sequences from human intestinal spirochaetosis. Isolates in red are sequenced in this study, the ones in blue are from ^36^, isolates in green from ^35^, and those in orange from ^37^. The phylogenetic clusters are annotated according to the terminology used in the three papers respectively.

### The Brachyspira aalborgi species is highly divergent

Closer investigation and comparative genomic of the 15 *B. aalborgi* CS genomes show a relatively high heterogeneity between isolates. The average nucleotide identity (ANI) ranged between 97.07% and 99.93% (average 97.49%), with an average of 86.09% (76.4 - 100%) aligned percentage, see Fig. S1A, Table S6. This could be compared with the divergence between the 4 sequences available from *B. pilosicoli* isolates (98.14 - 98.82 %, average 98.32 %) and for the 24 publicly available whole-genome sequenced *B. hyodysenteriae* (98.92 - 99.96 %, average 99.13 %. Also, the dDDH values between the *B. aalborgi* strains were sometimes below the 70% percent that is the suggested threshold for a species, on average 73.6% (68.4 - 99.6%). The PC3714II isolate, which grouped away from the other *B. aalborgi* in the 16S analysis, did not show any deviant characteristics neither based on the *nox* analysis, ANI or dDDH.

### Genome content and comparative genomics

Annotation of the new genomes showed a predicted number of coding sequences (CDS) between 2294 and 2554 per genome (Table 3). Out of these, for on average 49 % (46 - 52 %) of the predicted CDS, no annotations of blast e-value < 1e^−9^ was found. To investigate the difference in functional potential between the isolates we performed pan-genome analysis using the Roary tool. The core genome size (number of genes present in >95% of the isolates) among the *B. aalborgi* isolates was 1498 out of the total pan genome of 4691 genes. This also revealed a large heterogeneity between strains with three clear clusters of strains with different gene content (Figure 4).

**Figure 4.**
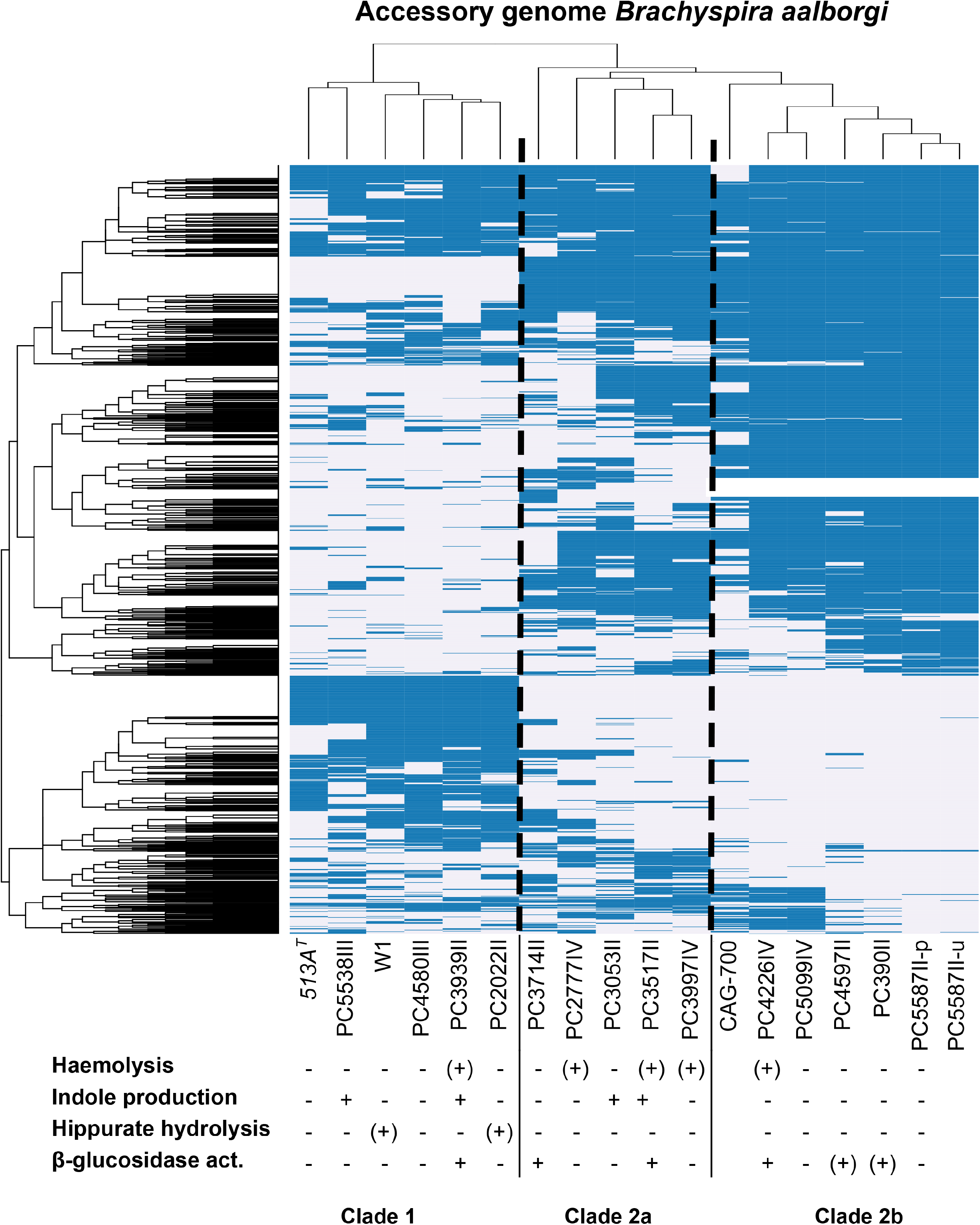
Heatmap showing hierarchical clustering of the accessory genome in *Brachyspira aalborgi, i.e*. genes present in < 99% of the genomes, together with the results from the biochemical characterization of isolates. Presence of an orthologous group is shown in blue and absence in white. Included genomes are the 16 newly sequenced *B. aalborgi* together with *B. sp*. CAG-700.

We also performed a pan-genome analysis including the previously described genomes of the *Brachyspira* genus. Using an amino acid identity cut-off of 70%, we identified 749 core genes that were common for over 99% of the genomes, out of a total pan genome of 8504 gene clusters. Out of the accessory genes 2026 were unique for strains of *B. aalborgi*, out of which 418 were common for all the 18 genomes. In the same comparison 1003 genes were only found in *B. pilosicoli* strains, out of which 425 were common for all the 5 *pilosicoli* genomes included in the comparison.

Using tools to identify bacterial secretion systems and CRISPR-Cas systems showed all isolates to have genes encoding for complete flagella systems but no other complete secretion systems. Using the prophage prediction tool ProPhet revealed no putative prophage sequences in any of our newly sequenced genomes.

### Host interaction factors

Further scrutiny of the *B. aalborgi* pan genome revealed several genes encoding for sialidase in the genomes, with a different number of genes in the different strains. Most extreme is the reference strain *513A^T^* where no sialidase gene was annotated, to PC3714II and PC4226IV that harboured 13 genes encoding for sialidase. Most of the predicted sialidase genes were annotated as sialidase *nedA* by homology to UniProtKB Q02834, a sialidase from *Micromonospora viridifaciens*, but also to UniProtKB P15698, a sialidase from *Paeniclostridium sordellii* and sialidase *nanH*, UniProtKB P10481 from *Clostridium perfringens* (Fig. S2). Interestingly, many of these genes were unique for *B. aalborgi* in the pan-genome analysis described above.

We also identified another gene encoding for a potential mucin-degrading enzyme in the core genome, namely the protease YdcP, which by COG was classified as a Collagenase-like protease, PrtC family. This gene was present in all other *Brachyspira* genomes studied, as well as another noteworthy core gene in *aalborgi*, encoding for TolC, an outer membrane protein required for the export of virulence proteins and toxic compounds without a periplasmic intermediate ^54^.

### Spirochaetosis is not detected using standard primers for GI tract microbiota analysis

To verify the presence of *Brachyspira* bacteria in the colonic microbiota using molecular techniques we used 16S amplicon sequencing where the 16S ribosomal RNA was amplified from DNA extracted from the sigmoid biopsies. The method and primer combination used are among the most commonly used to study the human gastrointestinal microbiota and amplifies the V3-V4 region of the 16S rRNA. Examining the classification of the reads it was noteworthy that the level of OTUs classified to the *Brachyspira* genus and even to the Spirochaete phylum was very low (0 - 6.6 % of the reads, average 0.7 %) in the individuals with CS and completely undetected in the age- and sex matched controls (Fig. S3).

To investigate the amplification capacity we first looked into the potential for primer annealing bioinformatically and observed that the reverse primer had a mismatch in the non-degenerate positions close the 3’ end of the primer, which we suspected would lead to poor amplification efficiency (Fig. S4). To verify this experimentally we made mixes of genomic DNA from five of the *Brachyspira* isolates at a fixed DNA-to-DNA ratio (1:3) with a Microbial mock of known composition respectively to evaluate the performance of the primers. The results gave at hand that 0.05-0.7% of the amplified reads were classified as *Brachyspira*, much below the expected 25%. The higher number (0.7%) was seen for the *B. pilosicoli* strain PC5538III (Fig. S5).

Despite the failure of the primers to amplify *Brachyspira* 16S rRNA we evaluated whether CS status verified by histopathology was associated with alterations in the microbial community composition and diversity. However, this did not reveal any significant differences between the groups, as illustrated by the lack of segregation between the groups using principal component analysis (Fig. S6A), or in terms of Shannon diversity (p = 0.07) or Chao1 index (p = 0.41) (Fig. S6B and S6C).

### Looking for CS in the Human Microbiome Project data

To look for spirochaetosis in a publicly available dataset from another population-based microbiota study we downloaded the 16S amplicon data from stool samples from 324 individuals from the Human Microbiome Project (HMP) ^55^. These samples had been amplified using two different primer combinations compared to our dataset, one amplifying the V1-V3 region of the 16S rRNA and the other the V3-V5 region. We also downloaded assembled shotgun metagenome sequences from 179 individuals to see if we could find any signs of *Brachyspira* sequences in that data. We found two shotgun metagenome assemblies containing *Brachyspira* bacteria and those two samples also had 16S amplicon reads classified to the *Brachyspira* genus, however the levels were almost undetectable, 0.03 % (V3-V5) and 0.04 % (V1-V3) *Brachyspira* reads, respectively, (Fig. S7). These assemblies came from the same individual, albeit from different stool samples. Further investigation of the primer pairs used for the two amplification protocols used show that, alike 341F/805R, both these primer combinations are likely to be very inefficient in amplifying *Brachyspira* 16S rRNA due to primer mismatches (Fig. S4).

## Discussion

Human colonic spirochaetosis (CS) is a condition of striking histological appearance with bacteria adhering to the colonic epithelium ^1,9,56^. Despite this, the clinical impact of CS is not established and successful culture of spirochaetes from human material is rarely reported. Nor have any isolates from human colonic spirochaetosis (CS) been characterised at the whole genome level. In this study we report isolates and whole genome sequences from 14 individuals with CS, together with the *Brachyspira aalborgi* type strain *513A^T^* [ATCC 43994] and previously described clinical isolate W1 ^10^.

From 17 individuals with histological CS, isolation of spirochaete colonies from three of them failed. Interestingly, all three of these individuals were diagnosed with IBS based on Rome III criteria, and both of the individuals with IBS diarrhoea were among these. In total we only obtained viable isolates from 3 out of the 6 individuals with CS *and* IBS. Comparing the 16S rRNA sequences of the sequenced isolates with previously published studies of samples from human CS, isolates belonging to the cluster 2-4 in the phylogenetic tree are missing in our material. This could be because none of the individuals in our study carried these kinds of bacteria but could also be due to that only cluster 1 isolates are cultivable, as assumed by others ^35^. However isolates in cluster 1 also seem to be more common when other methods are used ^57^.

Trying to circumscribe the species of the isolates we were limited by the fact that no *B. aalborgi* isolates were hitherto whole genome sequenced, and we therefore also included the *B. aalborgi* type strain *513A^T^* in the sequencing. We then compared the 16S rRNA sequence of the newly sequenced isolates to publically available *Brachyspira* 16S sequences from the SILVA database as well as tried to delineate the phylogeny based on the NADH oxidase, *nox*, gene (Figure 2A and B) ^31^. This allowed us to classify 14 of the patient isolates to be *Brachyspira aalborgi* and one to be a *Brachyspira pilosicoli*, which was further confirmed with better resolution using ANI and dDDH comparison to publicly available *Brachyspira* whole-genome sequences (Fig. S1).

Using the comparative metrics described above, we could also see that the sequence diversity within *B. aalborgi* is high compared to the other *Brachyspira* species where several whole-genome sequences are available, despite that the CS isolates were all from a limited geographical area consisting of two adjacent parishes in Stockholm, Sweden. The proposed and generally accepted species boundary for ANI and dDDH values are 95-96 % and 70 %, respectively ^60^ and the lowest ANI values within the *aalborgi* genomes were 97.07 %. However, the lowest dDDH value was 68.4 %, and several strains showed this level of similarity to the type strain. This means that several isolates are on the borderline of qualifying as novel species but there is also noteworthy continuum of decreasing similarity values, and more isolates from human colonic spirochaetosis would be needed to correctly delineate further differentiation within the *Brachyspira* genus. Also, the biochemical tests did not show any correlation with the ANI, dDDH or accessory genome-based groupings of the isolates suggesting that these tests, which have been proven useful in characterization of several other *Brachyspira* species cannot be used for species designation of *B. aalborgi* isolates. A high sequence diversity and signs of extensive recombination have also been described in *B. pilosicoli* both using MLST ^61^, and in comparisons between the three previously sequenced *pilosicoli* genomes ^11^, but with the whole-genome sequences available to this date, this variation seems to be lower than within the *B. aalborgi* species.

To investigate whether the genomes encoded for proteins that could participate in host-bacterial interaction and/or explain the capacity of the spirochaetes to penetrate both the loose and inner, attached mucus layers in the colon, we searched the genomic content of the CS isolates for genes involved in mucus metabolism. We found an abundance of sialidases in the *B. aalborgi* genomes, ranging from 2-13 genes encoding for sialidases per isolate, with the one exception being the type strain *513A^T^* that interestingly did not harbour any sialidase genes. Sialidases are proteins that hydrolyse α2,3-, α2,6-, and α2,8- glycosidic linkages of terminal sialic acid residues in oligosaccharides, glycoproteins and glycolipids ^62^. Sialidated sugars are abundant in mucins and other glycoproteins on the intestinal epithelium where there is a gradient of increasing terminal sialic acids from the ileum to the rectum in humans ^63^. The purpose of this cleavage can be to release sialic acids for use as carbon and energy sources, whereas in pathogenic microorganisms, sialidases have been suggested to be pathogenic factors where the bacteria use the sialic acid to evade the immune system ^64^. No sialidase/sulfatase have been found in *B. hyodysenteriae* but in *B. pilosicoli* and *aalborgi* ^65^. Interestingly, the large majority of the sialidase genes identified in the *B. aalborgi* genomes did not have homologues in the other *Brachyspira* species according to the pan-genome analysis. This multitude of highly homologous genes also posed problems for the *de novo* assembly and several of the annotated sialidase genes were truncated due to contig breaks.

Another potential virulence factor in *Brachyspira aalborgi* is the collagenase PrtC. This protease was originally described in *Porphyromonas gingivalis* and has also been found in other human pathogens such as *Helicobacter pylori, Salmonella enterica, Escherichia coli*, and *Bacillus subtilis* ^66^. Bacterial collagenases have been highlighted in pathogenicity due to their ability to cleave extracellular matrix components and thereby facilitate colonization and invasion ^67^ and PrtC has also been suggested to act as an adhesin to collagenous structures ^68^.

Another gene encoding what seems to be a core protein in *Brachyspira* is TolC, located between the genes encoding for the AcrAB multidrug efflux pump system subunits AcrB and AcrA (membrane-fusion protein) ^69^. The AcrAB–TolC efflux pump, which spans the inner and outer membranes of the bacterium, is able to transport a broad range of structurally unrelated small molecules/drugs out of certain Gram-negative bacteria, and this AcrAB-TolC complex have been shown to confer antibiotic resistance and survival in the gastrointestinal tract ^70^ and have also been described in *B. pilosicoli* ^11^.

Detection of spirochaetosis by histology requires invasive methods such as colonoscopy and collection of biopsied for histology assessment. To see if the spirochaetes could be detected by standard 16S amplicon studies of the microbiota we performed such an analysis using a primer combination commonly used in population-based studies of the human microbiota. To our surprise, the level of reads classified to the *Brachyspira* genus was very low despite being amplified from a tissue locations adjacent to those where an abundance of spirochaetes could be observed under the microscope. To identify the reason for this detection failure we compared the primers used in our study and the HMP study, from which we also analysed samples, and which is also a large population-based study. The results are shown in Fig. S4 and verified that none of the three primer combinations can efficiently amplify *Brachyspira* 16S rRNA, in our study due to a mismatch in the 805R primer. The HMP studies either used V3-V5 357F + 926R or V1-V3 27F + 534R ^71^, for which V1-V3 amplification of *Brachyspira* is hampered due to a mismatch in 27F and for the V3-V5 amplification a mismatch in 926R. Considering that spirochaetosis based on observations in the PopCol study increase the risk for having IBS three-fold ^14^, this failure to detect *Brachyspira* using standard protocols for microbiota studies should be taken seriously since this potentially important etiological agent in the pathology of IBS will be missed.

The current study had a number of strengths being population-based with good representation of the general population ^17^. There were also weaknesses. We only obtained viable isolates from few individuals with IBS and considering this together with the large genetic heterogeneity between isolates we did not have statistical power to further investigate associations between bacterial genetic traits and lower gastrointestinal symptoms, including inflammatory parameters such as eosinophil infiltration. However, the notion of the poor success rate in culturing IBS isolates could also be important and should be investigated further. Also, studying more isolates and genomic sequences in the future from human CS in general would allow us to make more thorough inferences on functional genomics and pathogenic potential. Albeit looking similar morphologically in the tissue sections, when culturing the isolates we observed differences in phenotype, both in cell size, shape and growth rate. Previous studies have shown that there might exist a considerable complexity in the spirochaete population in each individual ^36^ but our approach of purifying and sequencing one isolate per individual did not allow us to investigate this. However, from one of the individuals we had two morphologically different isolates identified and from one the genome sequences turned out to consist of two species, suggesting there is more to be investigated in this aspect.

In conclusion, we report for the first time the whole genome analysis of a collection of human clinical isolates from individuals with colonic spirochaetosis where we show that genomic diversity between isolates is high. From one individual we could retrieve whole-genome sequences from two distinct species, confirming the observation of mixed infections. The sequencing of the *B. aalborgi* type strain also allowed us to make genome-based species circumscription with high resolution compared to previous 16S and *nox*-based investigations. We found several genes in the spirochaetosis genomes that suggest intimate host-microbe interaction, and that could confer a pathogenic potential to spirochaetes in the human colon, such as a multitude of genes encoding for mucin-degrading proteins. The association of these bacteria with for example irritable bowel syndrome and the notion that they go undetected in major studies of the gut microbiota raise questions as to whether they represent an underestimated etiological agent in functional bowel disorders, an issue that should be considered when designing future microbiota analyses.

## Acknowledgements

KT was supported by the Carl Lenhoff foundation for Spirochaetosis research. AA was supported by Ruth and Richard Julin Foundation and Nanna Svartz Foundation. The study was also supported by grants from the Söderberg Foundation and Medical Research Foundation to LE.

## Author contributions

KT performed the whole genome sequencing and data analysis, bioinformatics analyses, interpretation of the results, and wrote the manuscript. LI performed the biochemical characterisation of the spirochaetes. AB supervised the experimental work with the bacteria. ÅÖ and MMW performed the histopathology. LE, LA and NJT designed the PopCol study, and LA was principal investigator and responsible for the management of the study. LK led the data collection. AA characterised cases based on anamnesis and questionnaire data and participated in drafting the manuscript. LE was responsible for the management of the culturing and sequencing part of the study.

All authors critically revised the manuscript for intellectual content and approved the final version of the manuscript including the author list.

## Data availability

The genomes are submitted to GenBank under BioProject PRJNA513011.

---

***Figure S1***. Comparative genomics within the *Brachyspira* genus. A) Average Nucleotide Identity (ANI) comparison. B) Digital DNA-DNA Hybridisation (dDDH) comparison. C) Tree based on accessory genome analysis (presence/absence of accessory genes).

***Figure S2***. Maximum Likelihood tree of the genes annotated to encode for Sialidase in the *B. aalborgi* genomes. Leaves are coloured according to the source of annotation inference and the table is summarizing the number of sialidase genes per isolate.

***Figure S3***. 16S amplicon-based microbiota analysis in spirochaetosis cases and unaffected controls. A) Relative phylum composition in cases and controls. B) Proportion of reads assigned to the *Brachyspira* genus, cases and controls.

***Figure S4***. Primer mismatches in the different primer combinations compared to *Brachyspira* 16S rRNA sequences. Magnified areas show WebLogo representation of the *Brachyspira* 16S alignment in the primer target regions, together with the primers in question. Boxes mark the bases with mismatches.

***Figure S5***. Verification of primer failure in samples of known compositions. All samples were containing a 1:3 proportion of pure *Brachyspira* DNA and a mock community. A) Phylum composition in the different samples. B) Proportion of reads classified to the *Brachyspira* genus depending on the *Brachyspira* species spiked in. The left-most column represents the expected distribution.

***Figure S6***. 16S amplicon-based microbiota analysis in spirochaetosis cases and unaffected controls. A) Principle component analysis of OTU composition in cases and controls. B) Shannon & Chao1 diversity

***Figure S7***. Proportion of reads assigned to the *Brachyspira* genus in 324 individuals from the Human Microbiome Project (HMP).

